# Mechanical evolution of 3T3 fibroblastic cells exposed to nanovibrational stimulation

**DOI:** 10.64898/2026.04.09.717227

**Authors:** O. Johnson-Love, F. M. Espinosa, J. R. Tejedor, G. Gorgone, P. Campsie, M. J. Dalby, S. Reid, R. Garcia, P. G. Childs

## Abstract

Cells are mechanosensitive, responding to external mechanical stimulation. Nanovibrational stimulation has been shown to enhance cell contractility and actin stress fibre formation. These changes in morphology occur quickly, alongside associated mechanical changes. Here, the relationship between acute morphological and mechanical changes in NIH 3T3 fibroblastic cells in response to nanovibrational stimulation is presented. A 1 kHz, 30 nm vibration is applied continuously for 72 hours. Atomic force microscopy (AFM) quantifies mechanical properties of the nucleus and cytoplasm at multiple timepoints, while immunofluorescence tracks morphological changes. Within 3 hours of stimulation, both nuclear and cytoplasmic stiffness increase significantly, accompanied by a decrease in the cellular fluid exponent, suggesting a shift of the cell towards more solid-like behaviour. These changes correlate with increased nuclear area. Actin polymerisation also increases within 24 hours, although variably. To understand the role of the cytoskeleton, actin polymerisation and contraction are inhibited using cytochalasin D and blebbistatin. Results show that inhibition prevents stiffness increases and results in a higher fluid exponent, indicating a more fluid-like state. These findings demonstrate that actin-myosin dynamics mediate cell stiffening under nanovibrational stimulation. Interestingly, prolonged stimulation appears to reverse this effect, suggesting that temporal optimisation of stimulation may enhance long-term mechanotransducive responses.

## 1. Introduction

Cells are intrinsically mechanosensitive, continuously responding to forces from their surrounding environment through structural and biochemical changes [1-3]. Mechanical cues are sensed via mechanosensitive ion channels and transmitted through integrins to intracellular adapter proteins such as talin [4, 5]. Force-induced unfolding of talin exposes vinculin binding sites and triggers downstream signalling and cytoskeletal reorganisation [5, 6]. In response to these external forces, cells can migrate, differentiate, and adapt their shape and configuration [7]. Cell mechanics are governed by the cytoskeleton, a dynamic network of filaments and crosslinking proteins that extends throughout the cell [8]. Actin filaments mediate adhesion, motility and cytoskeletal tension, with contractions generated through interactions with the motor protein myosin [9-11]. Actin-myosin networks link focal adhesions at the cell surface to the nuclear envelope, forming a continuous mechanical pathway in which forces are transmitted via retrograde actin flow and the molecular clutch mechanism [12, 13].

Mechanical forces transmitted to the nucleus can deform its structure, reorganise chromatin and influence epigenetic markers, leading to altered transcriptional activity [14-19]. Such nuclear mechanotransduction directly affects gene expression and contributes to cell fate decisions and lineage commitment [20]. Externally applied mechanical cues such as vibrational stimulation offer a powerful means of modulating mechanotransduction.

Nanometer-scale vibrational stimulation has been shown to modulate cellular behaviour by targeting mechanotransductive components such as integrins, adhesion complexes, and mechanosensitive ion channels including Piezo2, TRPV1 [21, 22]. This stimulation has shown effects in a number of cell types, including promotion of stem cell osteogenesis, bacterial biofilm inhibition, fibroblast fibrosis and monocyte differentiation [21, 23-25]. Justification for nanoscale vibration can be found from several cellular properties. For example, within focal adhesions, talin is a critical mechanosensor which unfolds in 30-50 nm increments, exposing binding sites for vinculin that activate downstream signalling cascades [6, 26, 27]. Additionally, nanoscale undulations of the cell membrane during adhesion and migration may influence how cells sense and respond to their environment [28]. Despite these anticipated changes, the effects of nanovibration on cellular stiffness, and the relationship between mechanical and morphological changes, have not been investigated.

To address this gap, stiffness measurements require sensitive techniques capable of capturing dynamic changes in cell mechanics. Atomic force microscopy (AFM) is a widely used technique for characterising cell mechanics, including stiffness and viscoelastic properties [29, 30]. Previous AFM studies have demonstrated that mechanical loads such as shear stress, point loading and tensile strain, increase cellular stiffness and induce cytoskeletal reorganisation [31-33]. For example, MC3T3-E1 cells exposed to flow-induced shear stress stiffened 1.7-fold compared to static controls, correlating with actin rearrangement that persisted after flow cessation [31]. Static point loading increased cell stiffness by 1.35-fold and induced actin reorganisation, whilst 4% uniaxial stretching of 3T3 fibroblasts produced a measurable stiffening response [32, 33]. Together, these studies show that applied forces can induce lasting changes in cell stiffness through cytoskeletal adaptation.

Living cells exhibit viscoelastic behaviour, therefore measuring both elastic and viscous components is essential [34]. However, AFM studies often rely on purely elastic models such as Hertz or Sneddon, which may oversimplify cell mechanics [35, 36]. The molecular clutch mechanism highlights the importance of viscoelasticity, as clutch engagement depends on both substrate stiffness and viscosity [13, 37]. On sufficiently stiff or viscous substrates, retrograde actin flow slows, allowing talin to unfold and reinforce adhesion [13, 37]. In addition, cells are typically cultured on rigid surfaces in the GPa range; finite cell thickness introduces substrate effects that can distort AFM measurements [30]. Garcia and colleagues have developed methods to correct for these artefacts and extract viscoelastic parameters using a power-law model, enabling the separation of elastic and viscous components [30, 38]. The modulus of the cell can be described by the following expression,

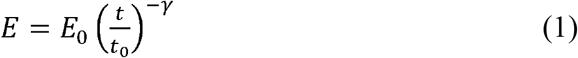

Where E_0_ is the compressive modulus of the material (or cell) at time t_0_ whilst γ is the fluidity exponent. When γ = 0, the material can be considered elastic, whilst γ = 1 defines a Newtonian liquid [38].

Whilst studies have investigated how cells respond to different loading conditions, mostly within the μm – mm range, the specific stiffening response to nanomechanical stimulation remains largely unexplored [39]. Furthermore, temporal changes in stiffness are rarely examined. Here, this study addresses this gap using NIH 3T3 fibroblasts, which are central to connective tissue maintenance and known to be mechanoresponsive [40-45]. In this work, 1 kHz, 30 nm vibration was applied to cells with AFM and fluorescence microscopy used to characterise mechanical and morphological responses. It was hypothesised that nanovibrational stimulation would induce cytoskeletal reorganisation, producing measurable and dynamic changes in cell stiffness.

## 2. Results and Discussion

### 2.1 Nanovibration alters cytoskeletal organisation, adhesion formation and nuclear morphology

Nanovibrational stimulation of 1 kHz, 30 nm was applied for 72 hours with cell response assessed within this timeframe. Measurements were obtained from nanovibrated (NV) and control (CL) cells after 3, 24, 48 and 72 hours of stimulation or static culture. Nuclear area in vibrated cells was found to increase within the first 3 hours of stimulation as shown in Figure 1A. Although this response was not sustained for the entirety of stimulation, another significant increase was seen after 48 hours of stimulation. Cell area was increased by nanovibration after 24 hours, however at later timepoints this was no longer significant (Figure 1B). Actin staining intensity, relating to actin polymerisation, was also found to be significantly increased in nanovibrated samples within both the first 3 and 24 hours of stimulation (Figure 1C). Similar to nuclear area, this response did not remain raised for the entirety of stimulation. Vinculin expression (total fluorescent intensity of vinculin per cell) was also shown to increase within the first 3 hours of stimulation and again after 72 hours of stimulation although there were no significant differences at other timepoints (Figure 1D).

**Figure 1:**
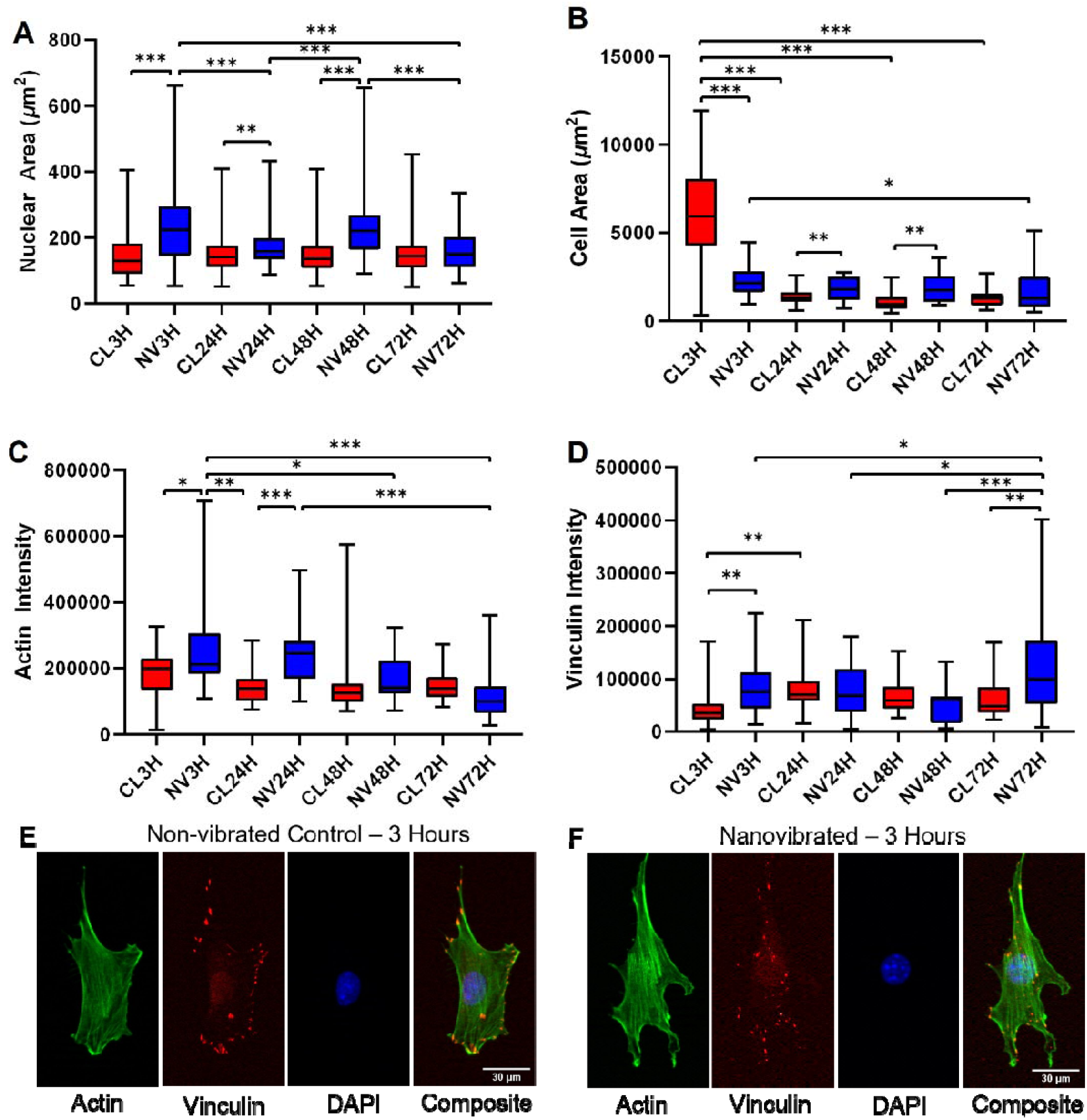
Morphology data of nanovibrated and control cells at multiple timepoints during nanovibrational stimulation. A) Nuclear area of cells across 72 hours of stimulation (Number of cells per sample: N > 100). B) Cell area of cells across 72 hours of stimulation (N ≈ 30). C) Actin intensity of cells across 72 hours of stimulation (N ≈ 30). D) Vinculin intensity of cells across 72 hours of stimulation (N ≈ 30). E) Immunofluorescent images of non-vibrated control, 3 hours after vibration began on vibrated samples. F) Immunofluorescent images of nanovibrate samples following 3 hours of vibration. Boxes denote interquartile range and whiskers show maximum and minimum values, significance was demonstrated by p < 0.05 (*p < 0.05, **p < 0.01, ***p < 0.001).

The early increase in nuclear area may be due to the cell flattening in response to vibration as the cell anchors itself more firmly to the substrate. Vinculin expression data appears to support this, showing an increase within the first 3 hours of stimulation, suggesting higher focal adhesion formation. An increase in actin polymerisation may also be suggesting increased actin tension, with the cytoskeleton pulling on the nucleus, leading to an increase in nuclear area. With this, the nuclear membrane may be mechanically stretched.

### 2.2 Cellular stiffness peaks earlier when vibrationally stimulated

AFM (JPK system) was used to measure mechanical changes in the nucleus and cytoplasm at the same time points that morphological changes were studied. Sequential measurements on cells were obtained from the same dish prior following 3, 24, 48 and 72 hours of stimulation. A non-vibrated control sample was also measured at the same timepoints, as well as prior to stimulation being applied (0H).

AFM measurements found the stiffness of the nucleus and cytoplasm to increase significantly after 3 hours of stimulation compared to non-vibrated controls (Figure 2A/Figure 3A and Figure 2B/Figure 3B). Whilst this stiffness reduced at later timepoints, the nucleus of nanovibrated samples remained stiffer than non-vibrated controls at 72 hours. The fluidity exponent of the nucleus did not show consistent changes between nanovibrated and non-vibrated controls at the same timepoints (Figure 2C/Figure 3C), however for the cytoplasm, the fluidity exponent was significantly reduced due to vibration, at all timepoints except 72 hours (Figure 2D/Figure 3D). Crucially, we also see a peak in stiffness for the control cells (nucleus and cytoplasm) but this is delayed to the 24 hour timepoint. This suggests that vibration induced stiffening may be related to acceleration of changes seen in the control cells.

**Figure 2:**
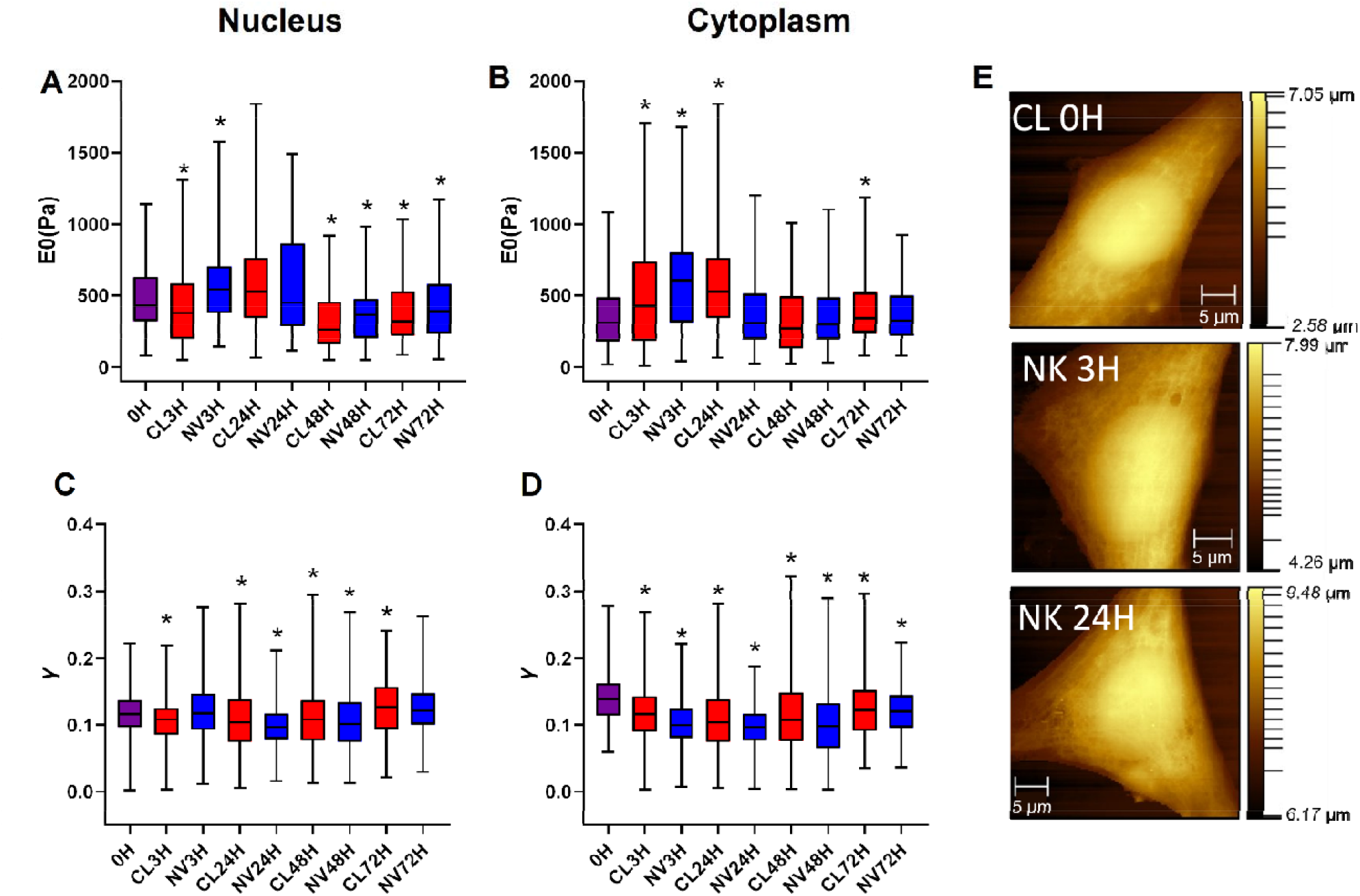
AFM measurements of NIH 3T3 cells at multiple timepoints during nanovibrational stimulation using a JPK AFM with corresponding control cell measurements. A) Compressive modulus (stiffness) of the nucleus shown to increase in nanovibrated samples after 3 hours compared to 0 hour timepoint controls. B) Similarly, the cytoplasm also shows an increased stiffness in nanovibrated samples after 3 hours compared to 0 hour timepoint controls. C) Fluidity exponent following stimulation in the nucleus of the cell. D) In the cytoplasm, fluidity exponent shown to initially decrease in nanovibrated cells. E) Topography images of non-vibrated control cell and nanovibrated cells following 3 and 24 hours of stimulation. Boxes denote interquartile range and whiskers show maximum an minimum values. Asterisks indicate statistically significant differences between sample and the 0H condition (p < 0.05). (n=10 cells, 6 FDCs per nucleus/cytoplasm).

**Figure 3:**
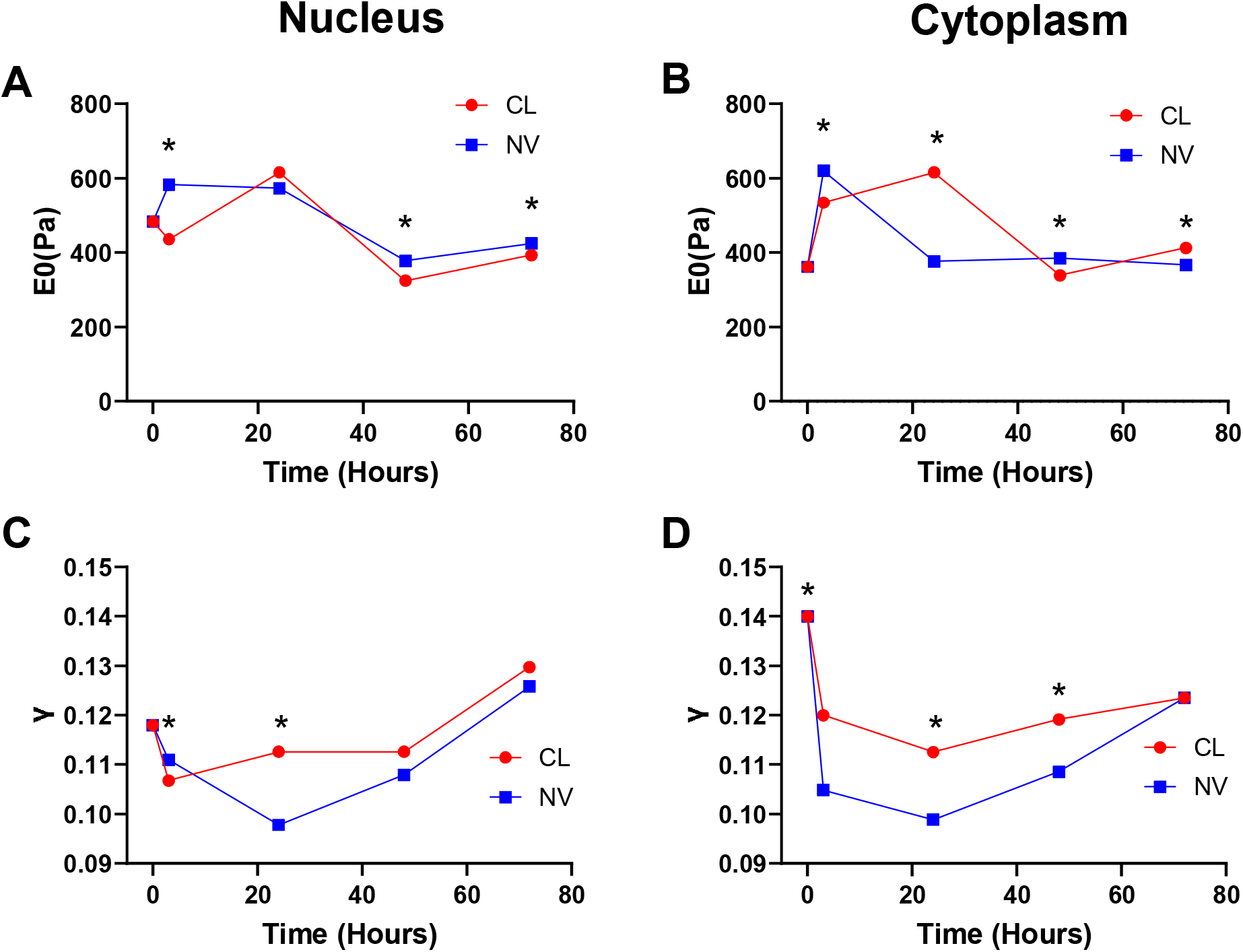
AFM measurements of NIH 3T3 cells at multiple timepoints during nanovibrational stimulation using a JPK AFM with corresponding control cell measurements. A) Compressive modulus (stiffness) of the nucleus shown to increase in nanovibrated samples compared to non-vibrated controls. B) Similarly, the cytoplasm also shows an increased stiffness in nanovibrated samples compared to non-vibrated controls C) Fluidity exponent following stimulation in the nucleus of the cell. D) In the cytoplasm, fluidity exponent shown to initially decrease in nanovibrated cells. Asterisks denote statistically significant differences between groups at that timepoint (p < 0.05). Standard deviation bars have been omitted for clarity (n=10 cells, 6 FDCs per nucleus/cytoplasm). 0H measurements were taken from control dish.

Further AFM measurements were taken using an Asylum AFM (Figure 4). Here, measurements were taken prior to nanovibrational stimulation being applied (NV0H) and, in the same dish, after 3, 24, 48 and 72 hours. A strong stiffening response was again observed following 3 hours of stimulation in both the nucleus and the cytoplasm (Figure 4A and Figure 4B). Again, this stiffness was not maintained and decayed over time showing a stronger trend than the previous experiment (Figure 2A/Figure 3A and Figure 2B/Figure 3B). The fluidity exponent decreased within the first 24 hours of stimulation, before gradually increasing at later timepoints (Figure 4C and Figure 4D). The trend between the compressive modulus and the fluidity exponent again support that the nucleus and cytoplasm are stiffening in response to vibration whilst simultaneously becoming less viscous. With increased duration of stimulation, the stiffness decreases whilst the cell becomes more fluid-like.

**Figure 4:**
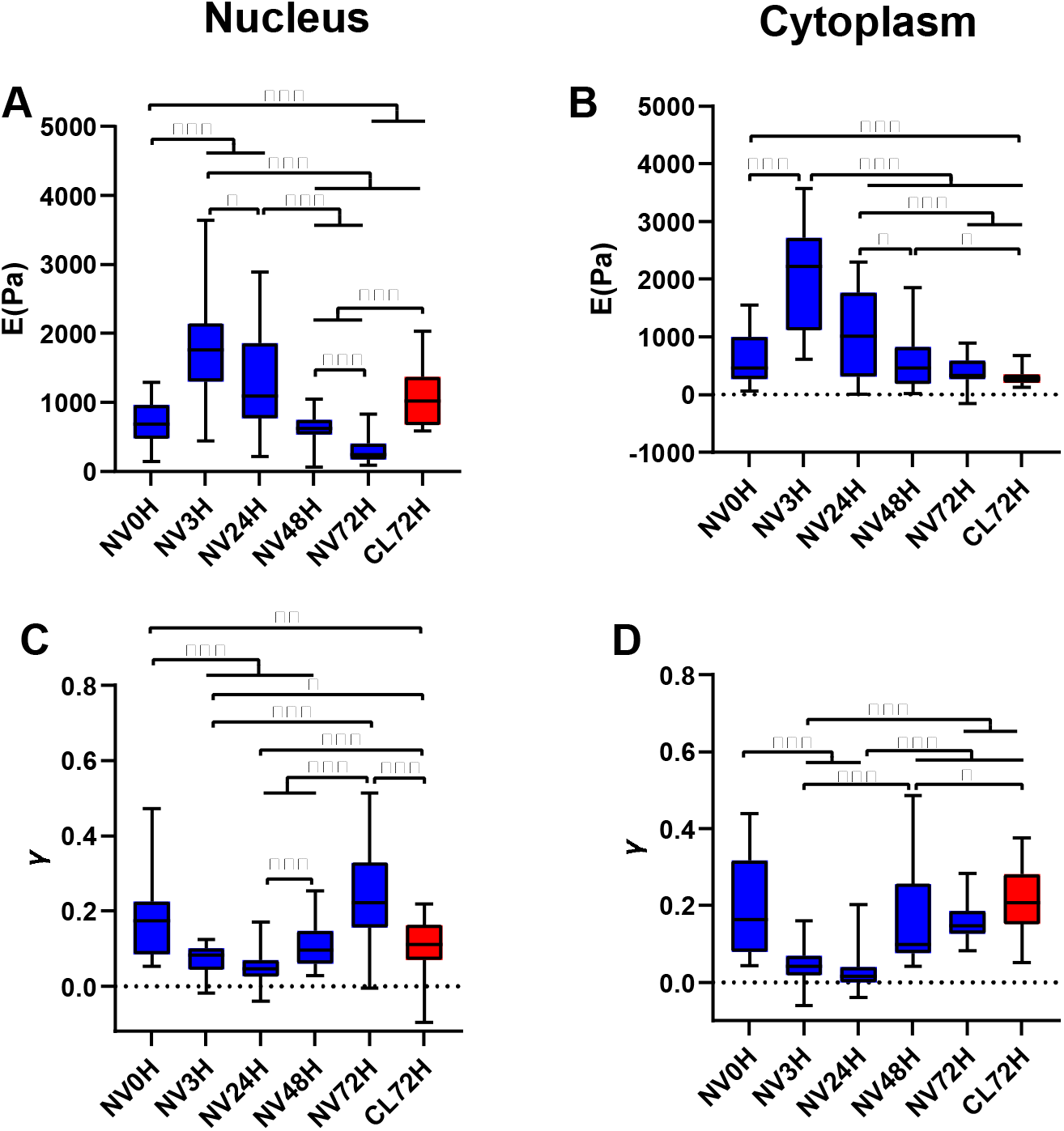
AFM measurements of NIH 3T3 cells at multiple timepoints during nanovibrational stimulation using an Asylum AFM. A) Nuclear stiffness shown to increase significantly in the first 3 hours of stimulation before decreasing at later time points. B) Cytoplasm stiffness shows a similar response, stiffening following 3 hours of stimulation and reducing at later time points. C) Fluidity exponent shows an initial decrease in the nucleus in the first 24 hours of stimulation before recovering at later time points. D) Cytoplasm again showed a similar response, decreasing within the first 24 hours before again recovering at later timepoints (n=10 cells). Boxes denote interquartile range and whiskers show maximum and minimum values, significance was demonstrated by p < 0.05 (*p < 0.05, **p < 0.01, ***p < 0.001).

Non-vibrated control cells measured at the 72-hour timepoint (CL72H) showed an increase in nuclear stiffness compared to measurements made prior to stimulation (NV0H). This may be due to the cells having an increased cell-to-cell contact at this later timepoint, representing a confounding factor and potentially resulting in mechanical changes. The increase in cellular stiffness due to vibration corresponds with the increase in both nuclear area and actin staining. The increase in nuclear area may be resulting in higher tension within the nucleus, and as a result a higher nuclear stiffness as was observed. An increase in actin polymerisation would be expected to coincide with an increase in overall cell stiffness and a decrease in the fluid exponent as the cell became more solid-like. To confirm this, and whether the actin cytoskeleton was responsible for the increased stiffness observed in the nucleus, actin contractility and polymerisation were subsequently inhibited.

### 2.3 Nanovibration induced cell stiffness is driven by actin polymerisation and contractility

Hypothesising that the changes in actin were driving the vibration induced stiffening, actin inhibitors blebbistatin and cytochalasin D were used to inhibit cell contractility and polymerisation respectively. AFM measurements were taken using an Asylum AFM at the same time points as previously as shown in Figure 5. Inhibitors were not applied until after measurements were taken at the 0H timepoint. As before, timepoint measurements were made to the same dish for each condition.

**Figure 5:**
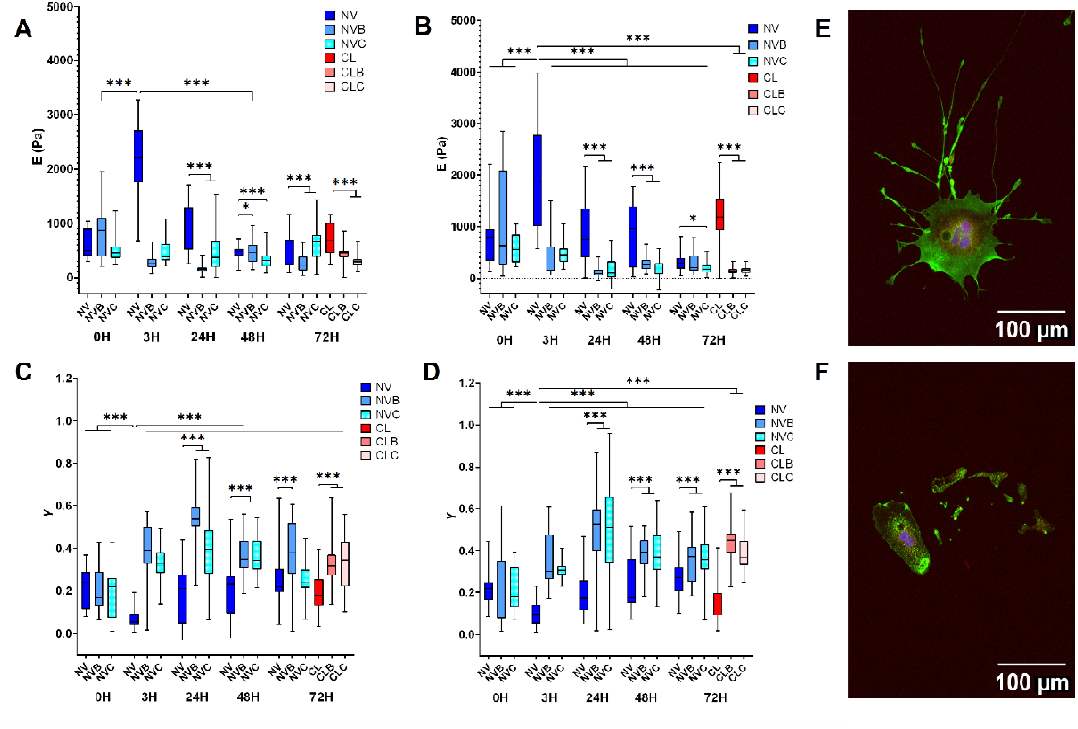
Mechanical data for actin inhibited cells. Blebbistatin was used to inhibit actin contractility whilst cytochalisin D was used to inhibit actin polymerisation. NV denotes nanovibrated, CL denotes control cells, B denotes addition of blebbistatin and C denotes addition of cytochalasin D. Inhibitors were not applied until after measurements were taken at the 0H timepoint. A) Application of both inhibitors resulted in a lower nuclear stiffness value following stimulation at all timepoints. B) Similar results were seen in the cytoplasm. C) Nuclear fluidity exponent saw higher values in inhibited cells following stimulation. D) Similar results were again seen in the cytoplasm (n=10 cells). E) Immunofluorescent image of cell exposed to blebbistatin with the actin stained in green, vinculin in red and DAPI in blue. F) Immunofluorescent image of cell exposed to cytochalasin D, again with actin stained in green, vinculin in red and DAPI in blue. Boxes denote interquartile range and whiskers show maximum and minimum values, significance was demonstrated by p < 0.05 (*p < 0.05, **p < 0.01, ***p < 0.001).

Inhibited cells were lower in stiffness in both the nucleus and cytoplasm in both nanovibrated and non-vibrated controls at all timepoints measured. Non-inhibited cells showed a similar stiffness response to previous experiments, with a significant increase in both nuclear and cytoplasm stiffness observed after 3 hours of stimulation. Later time points did not show a significant increase in stiffness in vibrated cell samples. The fluidity exponent was found to be significantly higher in inhibited cells in both the nucleus and cytoplasm.

Without actin contractility or polymerisation, cellular stiffness did not increase in response to nanovibrational stimulation. This suggests that actin fibre polymerisation increases following vibration, as observed in the increased actin staining intensity within the first few hours of stimulation. These actin fibres may then be pulling on the nucleus as the cell forms more focal adhesions with the substrate, as shown with an increased nuclear area and vinculin intensity. These morphological changes appear to lead to the increased stiffness response observed in both the nucleus and cytoplasm of the cell within the first few hours of stimulation, however none of the responses observed are maintained throughout applied stimulation. The higher fluidity exponent observed in inhibited samples corresponds with the lower stiffness values, indicating that disruption in actin polymerisation and contractility results in a softer, more fluid-like cell.

## 3. Conclusion

This study presents the first evidence of the interplay between morphological and mechanical changes in response to high frequency, nano-amplitude vibrational stimulation. Vibration of 30 nm amplitude at 1 kHz resulted in significant changes in both cell stiffness and the fluidity exponent within the first few hours of stimulation, as measured on two different AFM systems. It was noted that non-vibrated control cells also exhibited a time-dependant stiffening behaviour, but nanovibration accelerated this process. Mechanical changes corresponded with morphological changes in the nucleus and increased actin fibre formation. The cell appears to respond to vibrational stimulation initially through morphological changes and actin fibre reorganisation which lead to increased tension and thus increased cell stiffness. When actin polymerisation and contractility are inhibited, the cell no longer increases in stiffness in response to vibration and instead becomes softer and more fluid-like.

Other studies measuring mechanical changes in cells exposed to various loads, such as flow-induced shear stress, point-loading and tensile stress, observed an increase in cell stiffness in response to the applied stimulation [31-33]. Alongside an increase in stiffness, actin cytoskeletal reorganisation was also observed. Our results agree with these previous studies, showing an initial increase in cell stiffness, in both the nucleus and cytoplasm, which corresponded with an increase in actin intensity. Previous studies have also shown that the nucleus accounts for 69.7% of the overall stiffness in MSCs, with the nucleus retaining cell stiffness following applied vibration [50]. This change in stiffness following vibration corresponded with morphological changes which suggested increased tension within the cell through increased actin cytoskeleton formation.

Until now, there have been no studies which have shown the mechanical evolution of vibrated cells over time. In this study, we have shown that vibrated cells respond initially to stimulation with an increase in stiffness, however this effect is not retained during continuous stimulation. This may suggest that temporal control of the vibration is required to maintain a mechanical response in cells. A number of studies have sought to use vibration to drive osteogenesis[39] and it has separately been shown that an increased cellular stiffness is linked to an increased osteogenic response in MSCs [51]. Future studies may therefore benefit from consideration of periodic, rather than continuous, stimulation to retain this increased cell stiffness and maximise related phenotypic changes.

## 4. Methods

### Cell Culture

Murine NIH 3T3 cells were cultured in Dulbecco’s modified essential medium (DMEM, Sigma), supplemented with 10% fetal bovine serum (FBS, Sigma) v/v, 1% minimum essential medium non-essential amino acid solution (MEM NEAA) v/v and 2% antibiotics v/v. Cells were then cultured within an incubator at 37°C with 5% CO_2_ and were passaged every 3-5 days. For experiments, cells were seeded at a density of 1 × 10^3^ cells/cm^2^ for immunofluorescent staining into 35 mm Petri dishes. Cells used were all under passage 30. To inhibit actin formation and contractility, cytochalasin B (14930-96-2, Sigma Aldrich) and blebbistatin (ab120425, Abcam) were used respectively. Inhibitors were added to cell media at a concentration of 5 μM for cytochalasin B and 50 μM for blebbistatin, and cell media was replaced with inhibited media 24 hours after seeding (0 hour timepoint).

### Nanovibrational Stimulation

To apply nanovibrational stimulation to cells, a bespoke device, based on piezo actuators, was used, applying vibration of 30 nm at 1 kHz frequency to cultureware. Development of the device was previously reported by Campsie *et al* [52]. The device was calibrated using a laser interferometer (SIOS Technik GmbH), ensuring that vibration was uniform across the surface. Briefly, cells were grown in petri dishes which were magnetically attached to the vibration surface. Stimulation was switched on at the 0 hour timepoint and ran continuously.

### Immunocytochemistry

Following 3, 24, 48 and 72 hours of vibration, cell media was removed and 4% Formalin solution (Sigma) was used to fix cells at room temperature for 15 minutes. The fixative was then removed and cells were washed with phosphate buffered saline (PBS) three times. 1 mL of a Triton X permeabilisation buffer was then added to cells for 5 minutes at room temperature. The permeabilisation buffer was then removed and cells were blocked for 5 minutes at room temperature using 1 mL of a 1% PBS/Bovine Serum Albumin (BSA) solution (w/v), to reduce non-specific antibody binding. Vinculin recombinant rabbit monoclonal antibody (AB_2532280, ThermoFisher Scientific) was added to cells at a concentration of 1:100 in 1% PBS/BSA (w/v) and cells were incubated at 37°C for one hour. The primary antibody was then removed, and cells were washed three times with 0.5% PBS/Tween (v/v). Following rinsing, secondary Goat Anti-Rabbit Texas Red antibody (4010-07, 2B Scientific) was added to cells at a concentration of 1:100 with 1% PBS/BSA (w/v) and cells were again incubated for one hour. Once the secondary antibody was removed, cells were again washed three times with PBS/Tween. Samples were then stained with Alexa Fluor 488 phalloidin (Thermo Fisher) diluted to a concentration of 1:100 with PBS for one hour at room temperature. Following staining, cells were rinsed once with PBS which was then removed and 100 µL of Invitrogen Fluoromount-G Mounting Medium with DAPI was placed on the cells.

### Microscopy and Analysis

Images were taken using a Zeiss microscope (Imager.Z1) at x20 magnification. Both laser intensity and exposure time were kept consistent between images of the same channel to ensure intensity results could be compared. Image analysis was then conducted in ImageJ. All samples were plated in triplicate and three multi-channel images were obtained per well (or Petri dish).

Nuclear area analysis was performed using thresholding, which provided an automated and quick method to measure nuclear area. The images were converted into 8-bit and the brightness/contrast adjusted before thresholding. Once thresholding had been performed, a watershed was performed on the image to ensure cells were separated before the analyze particle function was used to measure nuclear area. To analyse actin and vinculin intensity, only images of isolated cells were used in analysis (to minimise effects from cell-cell contact). Using thresholding and the analyse particle function, the outlines of single cells were defined using the actin channel. These outlines were then used on both actin and vinculin to obtain integrated density for each cell measured. To normalise against the background, the mean grey value of the background in each image was acquired in three empty regions of the image and the average was determined. To obtain the total intensity of actin and vinculin per cell, the following formula was used:□.

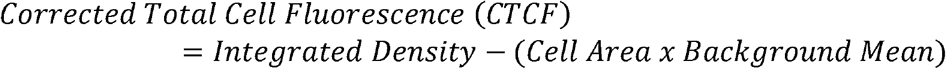

Intensity was plotted for both actin and vinculin staining. All results were plotted as box plots in OriginPro.

### Atomic Force Microscopy

A JPK Nanowizard 3 (JPK Instruments, Germany) and Asylum MFP-3D AFM were used to obtain FDCs. To obtain stiffness measurements, a silicon dioxide, spherical tipped cantilever (CP-PNPL-SiO-B) with a diameter of 3.5 µm was used. Cells were placed on a heated stage (37 °C) whilst obtaining measurements to ensure cells remained alive whilst AFM measurements were acquired. Once the system was calibrated, the AFM was set to contact mode. The nuclear and cytoplasmic stiffnesses of 10 cells were measured per condition. Due to additional automation, six points on the nucleus and six points on the cytoplasm of each cell were selected and measured on the JPK system, providing technical replicates. Force distance curves (FDC) were obtained, with measurements being taken prior to simulation beginning and after 3, 24, 48 and 72 hours of stimulation. Topography images were produced using a rectangular silicon tip on a silicon nitride cantilever (FASTSCAN-D_SS).

Elastic moduli and fluidity exponent values were extracted from the FDCs following the power law rheology model described by Garcia *et al*. previously[38]. Results were then plotted as a box plot to enable comparisons between samples.

### Statistics

Normality was firstly assessed using a Shapiro-Wilk test on the data. If normally distributed, a one-way ANOVA with a Tukey post-hoc test was used to compare the means of each experimental group. If not normally distributed, the non-parametric Kruskal-Wallis ANOVA with a Dunn’s post-hoc test was used. Any results with a p-value of less than 0.05 were deemed to be significantly different. A single asterisk was used in figures to denote p < 0.05, two asterisks denotes p < 0.01 and three asterisks denotes p < 0.001. All statistical analysis was performed in Origin Pro software.

## Acknowledgements

The authors acknowledge the financial support from EPSRC (EP/T517938/1, EP/X033554/1) and Scottish Funding Council. RG acknowledges funding by Comunidad de Madrid TEC-2024/TEC-158 (Tec4Nanobio-CM) and Horizon Europe MSCA Doctoral Network NANORAM, Grant No. 101120146).

